# Hydroxylation of antitubercular drug candidate, SQ109, by mycobacterial cytochrome P450

**DOI:** 10.1101/2020.08.27.269936

**Authors:** Sergey Bukhdruker, Tatsiana Varaksa, Irina Grabovec, Egor Marin, Polina Shabunya, Maria Kadukova, Sergei Grudinin, Anton Kavaleuski, Anastasiia Gusach, Andrei Gilep, Valentin Borshchevskiy, Natallia Strushkevich

**Affiliations:** Research Center for Molecular Mechanisms of Aging and Age-Related Diseases, Moscow Institute of Physics and Technology, Dolgoprudny, Russia; Institute of Bioorganic Chemistry, National Academy of Sciences of Belarus, Minsk, Belarus; Université Grenoble Alpes, CNRS, Inria, Grenoble INP, LJK, Grenoble, France; Institute of Biomedical Chemistry, Moscow, Russia; MT-Medicals LLC, Moscow, Russia; Institute of Biological Information Processing (IBI-7: Structural Biochemistry), Forschungszentrum Jülich, Jülich, Germany; JuStruct: Jülich Center for Structural Biology, Research Center Jülich, Jülich, Germany; Skolkovo Institute of Science and Technology, Moscow, Russia

**Keywords:** cytochrome P450, crystal structure, *Mycobacterium tuberculosis*, SQ109, CYP124

## Abstract

Spreading of the multidrug-resistant (MDR) strains of the deadliest pathogen *Mycobacterium tuberculosis* (Mtb) generates the need for new effective drugs. SQ109 showed activity against resistant Mtb and already advanced to Phase II/III clinical trials. Fast SQ109 degradation is attributed to the human liver Cytochrome P450s (CYPs). However, no information is available about interactions of the drug with Mtb CYPs. Here, we show that Mtb CYP124, previously assigned as a methyl-branched lipid monooxygenase, binds and hydroxylates SQ109 *in vitro*. A 1.25 Å-resolution crystal structure of the CYP124–SQ109 complex unambiguously shows two conformations of the drug, both positioned for hydroxylation of the ω-methyl group in the trans position. The hydroxylated SQ109 presumably forms stabilizing H-bonds with its target, the Mycobacterial membrane protein Large 3 (MmpL3). We anticipate that Mtb CYPs could function as analogs of drug-metabolizing human CYPs affecting pharmacokinetics and pharmacodynamics of antitubercular (anti-TB) drugs.

## Introduction

According to the World Health Organisation, Tuberculosis (TB) is one of the top 10 causes of death worldwide. Mtb is the leading cause of death from a single infectious agent [1]. Recently, multiple studies have reported the spread of MDR and extensively drug-resistant strains, which cannot be cured with first-line and even second-line anti-TB medications in the latter case [2,3]. The threat demands the development of novel drugs for anti-TB therapy, identification of their targets, and assessment of their metabolic stability.

One of the latest breakthroughs in this area was the discovery of SQ109 [4], which is currently in Phase II/III clinical trials for the treatment of MDR pulmonary TB [5]. SQ109 belongs to the 1,2-ethylenediamine class of anti-TB drugs [6], consisting of adamantane head and geranyl tail, proposed to disrupt the synthesis of the complex Mtb cell wall [7]. At least three mechanisms of action have been reported so far for the drug [8]. First, it inhibits transport of trehalose monomycolates by MmpL3 from the cytoplasm. Second, it inhibits respiration by blocking menaquinone biosynthesis by MenA and MenG. Finally, it acts as an uncoupler, collapsing the pH gradient and membrane potential. SQ109 has demonstrated promising inhibition of cell growth and a very low spontaneous drug resistance rates [6,7]. SQ109 showed *in vitro* activity against the known resistant Mtb strains [6] and *M. bovis* BCG [9]. It is also bactericidal against *M. smegmatis* (Msm) and *M. aurum*, although with reduced activity [9]. Moreover, SQ109 is active against non-mycobacterium species [10–13]. *In vivo* studies demonstrated SQ109 effectiveness in murine TB model [4]. SQ109 also interacts synergistically with other anti-TB drugs, such as rifampicin (RIF), isoniazid, and bedaquiline [9,14], which is crucial for the combination therapy [15].

SQ109 is effectively metabolized by human, dog, roten and murine liver microsomes [16]. Cytochromes P450 (CYPs), CYP2D6 and CYP2C19, were proposed to be primarily responsible for the metabolism in humans. However, to the best of our knowledge, no interactions with Mtb CYPs have been reported so far. Here, we show that Mtb CYP124 can bind and hydroxylate SQ109. CYP124 (gene Rv2266) has previously assigned activity towards methyl-branched lipids of isoprenoid origin [17–20]. Here we show, that SQ109 binds to CYP124 with Kd_app_ =3.4 ± 0.3 μM and is converted to the monohydroxylated product with a turnover number of 0.60 ± 0.09 min^−1^. We obtained 1.25 Å-resolution crystal structure, which revealed two close conformations of SQ109, confirming the formation of a ω-terminal hydroxy product. Finally, given the molecular docking, we propose that the newly formed hydroxy group of SQ109 affects its binding with a prospective drug target, i.e., MmpL3, stabilizing SQ109-OH in the binding pocket [21]. Our results support the hypothesis that SQ109 is a prodrug activated by Mtb CYPs. CYP124 represents the first example of a putative class of mycobacterial CYPs that might function similarly to drug-metabolizing human CYPs.

## Results

### CYP124, CYP125, and CYP142 bind SQ109

Multiple substrates have been previously identified for CYP124, including cholesterol, vitamin D, their derivatives [18,20], and methyl-branched lipids [17]. Although its physiological function (and, therefore, a natural substrate) is not yet confirmed, there is a common pattern among all the substrates – a methyl-branched tail, tightly positioned for ω-hydroxylation in the distal site. Therefore, we tested an anti-TB drug candidate SQ109 to be a CYP124 substrate, as its geranyl tail is well mimicking the 2,5-dimethylhexane group.

We performed a spectrophotometric titration assay to estimate the binding affinity of the drug to the active site. The UV-visible difference spectrum of CYP124 shows type I response upon addition of SQ109, with a characteristic peak at 386 nm and a trough at 421 nm, consistent with the known methyl-branched substrates [17,18,20] (Fig. 1a). The titration curve fits the tight-binding model with a Kd_app_ =3.4 ± 0.3 μM. The value is similar to that of for cholest-4-en-3-one (cholestenone) 3.4 ± 0.6 μM, and much stronger than that of for vitamin D3 (VD3) and 1α-hydroxy-vitamin D3 (1αOHVD3), 18 ± 2 μM and 34 ± 4 μM, respectively [20].

**Fig. 1.**
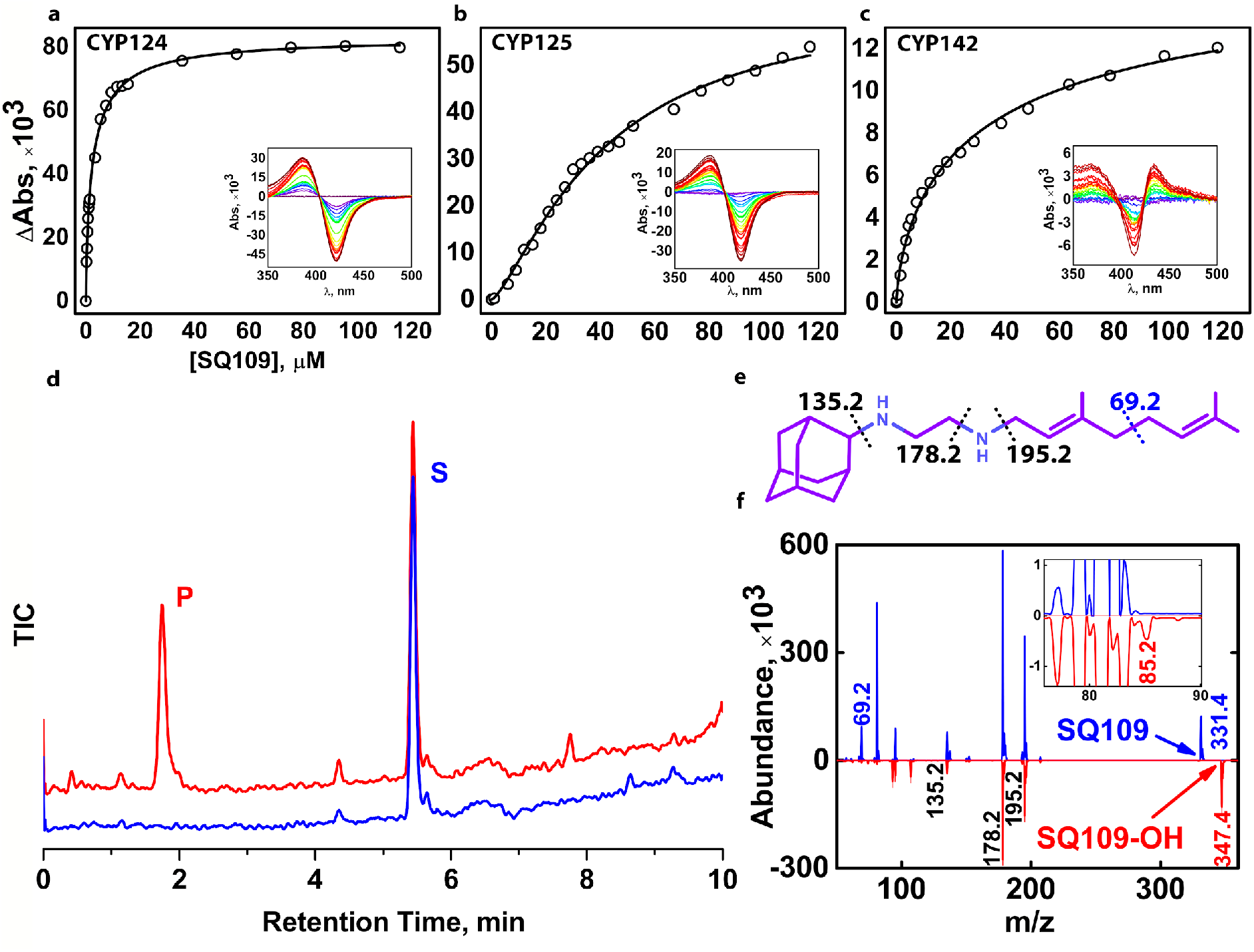
Binding and hydroxylation of a drug candidate SQ109 by Mtb CYPs. **a-c**, Difference spectra and titration curves of Mtb CYP124, CYP125, and CYP142 with SQ109, respectively. **d**, Formation of an SQ109 product by CYP124 in the reconstituted system, detected by HPLC. **e**, Structural formula of SQ109. Dotted lines show SQ109 fragmentation detected in the MS-MS spectra. **f**, MS-MS spectra of SQ109 (for parental molecular ion m/z = 331.4) is colored blue, and the product of SQ109 after CYP124 hydroxylation (for parental molecular ion m/z = 347.4) is colored red. The inset shows fragment ion m/z = 85.2 found only in the product spectrum and the corresponding to monooxygenated SQ109 tail moiety (original fragment ion of untreated SQ109 m/z = 69.2).

Besides CYP124, we also tested Mtb CYP125 and CYP142 for SQ109 binding. All these enzymes were previously proposed to have similar substrate specificity, as they were able to bind and hydroxylate cholesterol and its immunoactive derivatives [19,20]. Similarly to CYP124, CYP125 showed type I response upon addition of SQ109 (Fig. 1b). Surprisingly, SQ109 induced type II shift in the CYP142 spectrum, with a peak at 434 nm and trough at 414 nm (Fig. 1c), indicating an apparent interaction of the diamine group with the haem iron. Compared to CYP124, both CYP125 and CYP142 bind SQ109 with significantly lower affinity (Kd_app_ = 41 ± 3 μM and 52 ± 11 μM, respectively).

### CYP124 catalyses hydroxylation of SQ109

Next, we measured the catalytic activity in the reconstituted system, containing CYP124/125/142, SQ109, redox partners, and NADPH-regenerating system. LC-MS detected the putative product after an hour of incubation. Among three proteins, only CYP124 showed activity against SQ109 with a turnover number of 0.60 ± 0.09 min^−1^ (Fig. 1d). Although both CYP124 and CYP125 previously showed close substrate specificity [20], different stereometry of the binding [19] might explain the absence of the product for the latter. MS-MS subsequently identified the CYP124 product. The parental molecular ion m/z was 331.4 for the substrate and 347.4 for the product, suggesting the addition of one oxygen atom to SQ109 (Fig. 1f). The adamantane group can be easily identified by a characteristic peak at 135.2 (Fig. 1e). The peak at 69.2 in the substrate spectrum is of particular interest, as it corresponds to amylene and disappears in the product spectrum. In contrast, a peak at 85.2 appears in the product, indicating that CYP124 catalyzed the hydroxylation of SQ109 in the amylene part.

The previous study of *in vitro* metabolism of SQ109 by the MS analysis showed that SQ109 (m/z 331.5) was effectively transformed by microsomes from various species to four different products with m/z 361 (M1), 347 (M2), 195 (M3), and 363 (M4) [16]. Recombinant human CYPs, CYP2D6 and CYP2C19, could extensively metabolize SQ109 to M1 and M4, while CYP2C19 could also produce M2 and M3. Here, we show that Mtb CYP124 also contributes to the metabolism of SQ109 and catalyzes only the chemically disfavored ω-hydroxylation with m/z corresponding to M2 product.

### Crystal structure of CYP124–SQ109 complex

To determine the binding mode and the hydroxylation position, we co-crystallized SQ109 with CYP124 as described previously [20]. The structure of the CYP124–SQ109 complex was determined with 1.25 Å resolution (protein data bank accession number PDB ID: 6T0J). The overall structure resembles the CYP124–VD3 complex (PDB ID: 6T0G; ref. 20, Cα-Cα RMSD being 0.2 Å, Fig. 2a). 2*mF*_*o*_−*DF*_*c*_ composite omit map unambiguously shows SQ109 density in two conformations, which differ only in the ethylenediamine-adamantane part. At the same time, geranyl tails in both conformations are positioned at 4.0 Å above Fe in the typical geometry for ω-hydroxylation (Fig. 3a). It can be clearly defined that the carbon closest to the haem iron corresponds to the ω-methyl group in the trans position. Thus, CYP124 catalyzes the regio- and stereo-selective hydroxylation of SQ109.

**Fig. 2.**
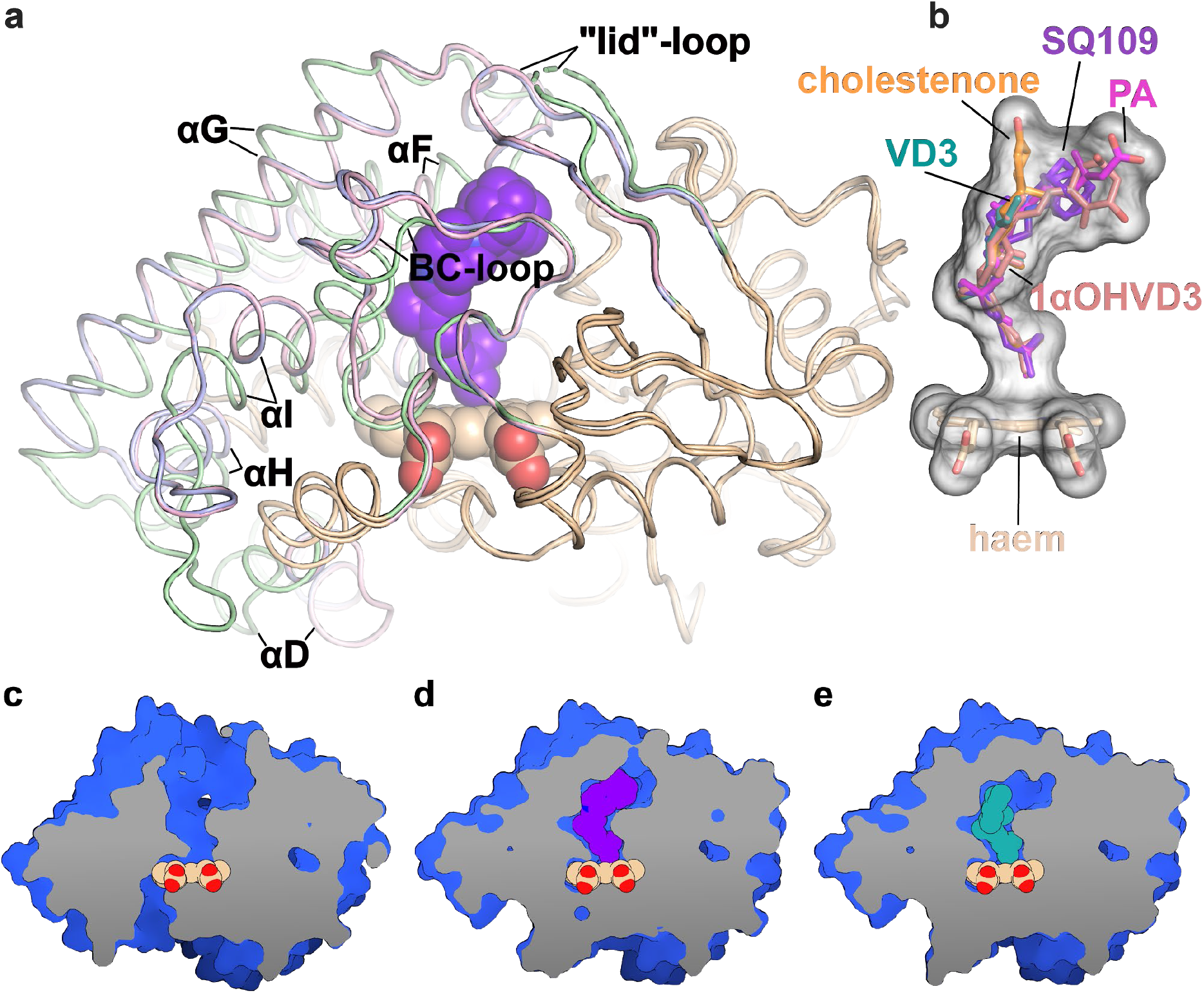
Structure of the CYP124 complex with substrates. **a**, Overlay of substrate-free CYP124 (PDB ID: 2WM5; ref. 17, pale green), CYP124–SQ109 (light pink) and CYP124–VD3 (PDB ID: 6T0G; ref. 20, light blue) structures. **b**, Superposition of SQ109 (purple), VD3 (PDB ID: 6T0G; ref. 20, green), 1αOHVD3 (PDB ID: 6T0H; ref. 20, pink), cholestenone (PDB ID: 6T0F; ref. 20, orange) and PA (PDB ID: 2WM4; ref. 17, magenta). **c-e**, CYP124 ligand-binding cavities for apo form, SQ109 and VD3 complexes, respectively.

**Fig. 3.**
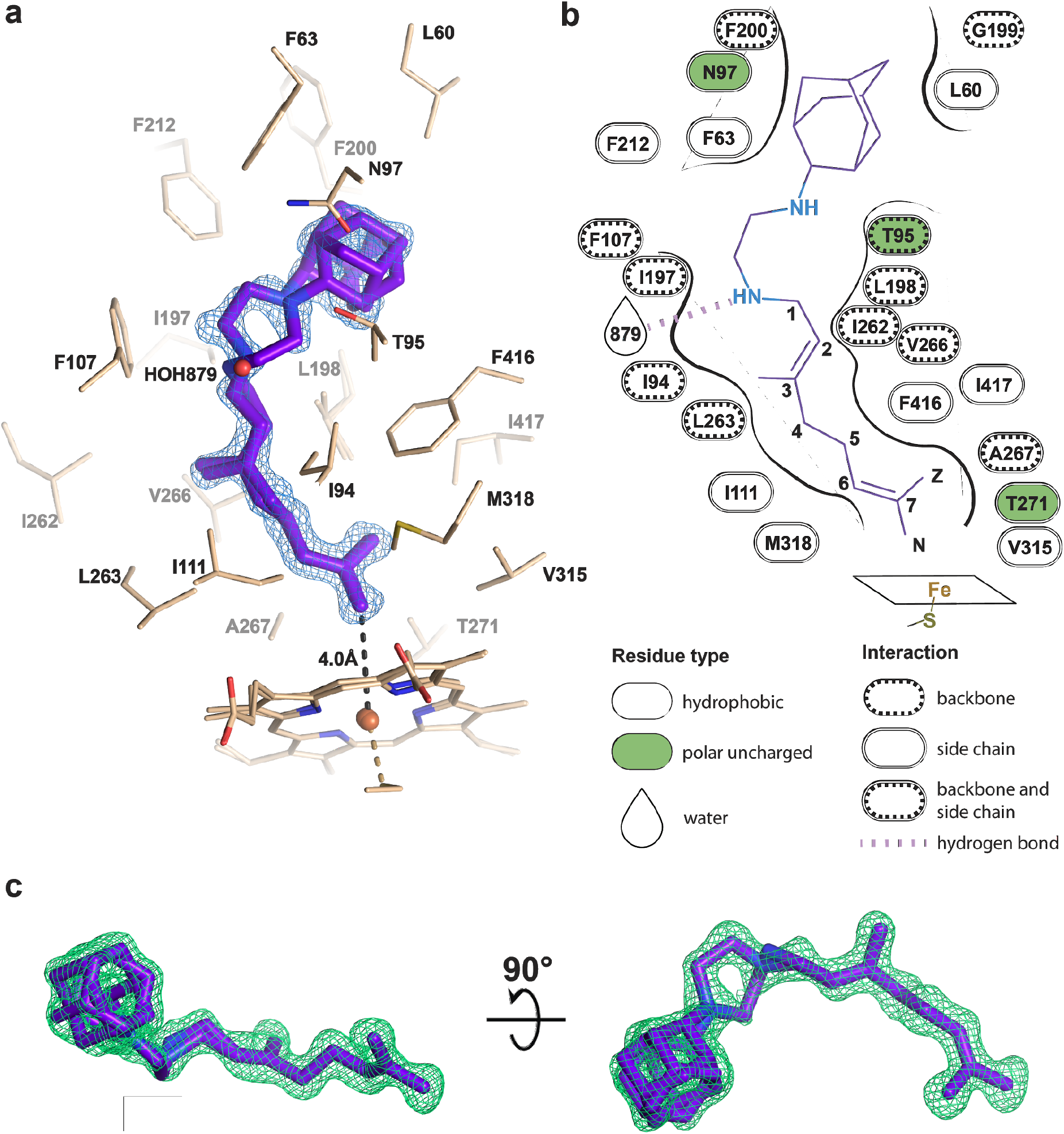
Binding pose of SQ109 to CYP124. **a**, CYP124 binding pocket with SQ109. Only side chains from the closest vicinity are shown for clarity. *2mF*_*o*_−*DF*_*c*_ composite omit map is contoured at 1σ. **b**, 2D diagram of the binding pocket. Weak H-bond with HOH879 (3.5 Å) is observed only for conformation A of SQ109 **c**, *mF*_*o*_−*DF*_*c*_ map contoured at 3σ in the final refinement step, which was used for SQ109 building.

Binding of SQ109 induces similar structural rearrangements, previously reported for phytanic acid (PA) complex and also observed for other substrates [17,20] (Fig. 2a). Briefly, paired movement of αF (K184–G199), BC-loop (S89-D115), and straightening of αI at N-terminal (S253-T271) results in the formation of a narrow tunnel, suitable for long methyl-branched hydrocarbons. The N-terminal of αD (K132–A139) is bent towards the proximal part of haem, contributing to the formation of the putative binding site for the redox-partner. Finally, FG-loop (G199–D208) and ordered “lid”-loop (W55-H69) enclose the active site from the solvent. The result of these transitions is shown for substrate-free, SQ109, and VD3 complexes (Fig. 2c-e, respectively).

The geranyl tail is tightly enclosed in the narrow tunnel with a broadening to accommodate ω-5 methyl branching position (methyl at C3 of SQ109, Fig. 3b). In contrast, adamantane and ethylenediamine are positioned more freely in a wide cavity in the upper part of the binding pocket. The interactions of SQ109 are mostly hydrophobic (L60, F63, T96, and F212 interact with the adamantane; F107, I111, L198, L263, V266, A267, F416, I417 fix the aliphatic chain), while in the predominant conformation (A), the drug could potentially form a weak hydrogen bond with HOH879 (Fig. 3, a and b). The structure of the complex again emphasizes remarkable flexibility, which allows CYP124 to accommodate a variety of chemically dissimilar substrates. Indeed, the superposition of all known substrates shows that the upper part of the active site can adjust to substantially different head radicals, while methyl-branched tail remains pinned for ω-hydroxylation (Fig. 2b; RMS distance between eight terminal carbons of the substrates not exceeding 0.6 Å).

Two common features between CYP124 substrates have been previously identified: a methyl-branched tail and a polar head [17]. The importance of the polar group remains unclear, as, in the majority of published complexes, it does not form H-bonds with the protein [17,20]. Moreover, in the CYP124–VD3 crystal structure (PDB ID: 6T0G; ref. 20), the polar A-ring is disordered in the upper part of the pocket (Fig. 2e). The absence of stabilizing H-bonds might result in the emergence of two conformations of the adamantane group. Although less polarity of the head group is not likely to deteriorate the CYP cycle, it may affect the reaction speed, as we have previously observed for VD3 and 1αOHVD3 [20].

## Discussion

The CYP family is well represented in Mtb, with 20 genes being identified while the function of the majority is still unknown. Their conservation during reductive genome evolution [22] indicates the importance of Mtb CYPs for survival or/and pathogenicity. Micromolar binding affinity and detected catalytic activity of CYP124 with SQ109, obtained in the study, indicate the CYPs’ potential role in the metabolism of xenobiotics in Mtb. Given the crystal structure of the CYP124–SQ109 complex and the functional assay, we identified that CYP124 hydroxylates SQ109 at the ω-methyl group in the trans position. The ability to hydroxylate SQ109 was not detected for the other two Mtb steroid-metabolizing CYP enzymes – CYP125 and CYP142, although they were both able to bind the drug.

SQ109 is a highly effective drug candidate against Mtb and, to a lesser extent, against Msm, with MIC values being 0.3-0.6 μM and 9.4 μM, respectively [8]. MmpL3 is suggested as one of the main targets of the compound in both organisms. Indeed, some mutations in the MmpL3 gene results in the emergence of the resistant strains [7,23]. The crystal structure of Msm MmpL3 in complex with SQ109 (PDB ID: 6AJG; ref. 24) showed that the drug disrupted core Asp-Tyr pairs (D256-Y646 and Y257-D645), apparently crucial for protein function [24]. However, SQ109 did not block the Mmpl3 flippase activity in spheroplasts, suggesting other molecular targets [25]. We consider a possibility that SQ109 is a prodrug, rather than a drug, which first needs to be activated by liver and/or Mtb CYPs. This assumption was first made by *Chen et al.* [9] based on the rapid compound metabolism by microsomal P450s [16]. The authors then hypothesized that the bactericidal activity of the drug might come from its metabolites, potentially produced by mycobacterial CYPs. They also noticed that the synergy with RIF [9,14] might be partly explained by the enhanced expression level of CYPs in RIF-treated mycobacteria [26].

To test this idea, we performed molecular docking of the determined metabolite – SQ109-OH, to the crystal structure of the Msm MmpL3-SQ109 complex (PBD ID: 6AJG) [24]. The structure shows that the newly formed OH group could fit within the SQ109 binding pocket and make favorable H-bonds (Fig. 4a). The top-ranked docking poses confirm the ability of SQ109-OH to H-bond with either S301 or the backbone oxygens of A637 and I297 (Fig. 4c). We also performed the molecular docking experiment using a generated homology-based model of Mtb MmpL3. Its binding pocket is somewhat similar to that of Msm, being different by only four residues, namely S301:A296, I319:T314, V638:L633, and L686:V681 (residue in Msm: residue in Mtb). The lost possibility of H-bonding with S301, in this case, might be compensated by bonding with S295 (Fig. 4e) or the Mtb-specific T314 (Fig. 4f), which was also confirmed by docking. The additional stabilization of SQ109-OH might facilitate the inhibition of the flippase activity observed for other Mmpl3-directed drugs [25], such as AU1235 [27] and BM212 [28]. Hence, our findings identify the first example of Mtb CYP participating in the biotransformation of anti-TB drugs.

**Fig. 4.**
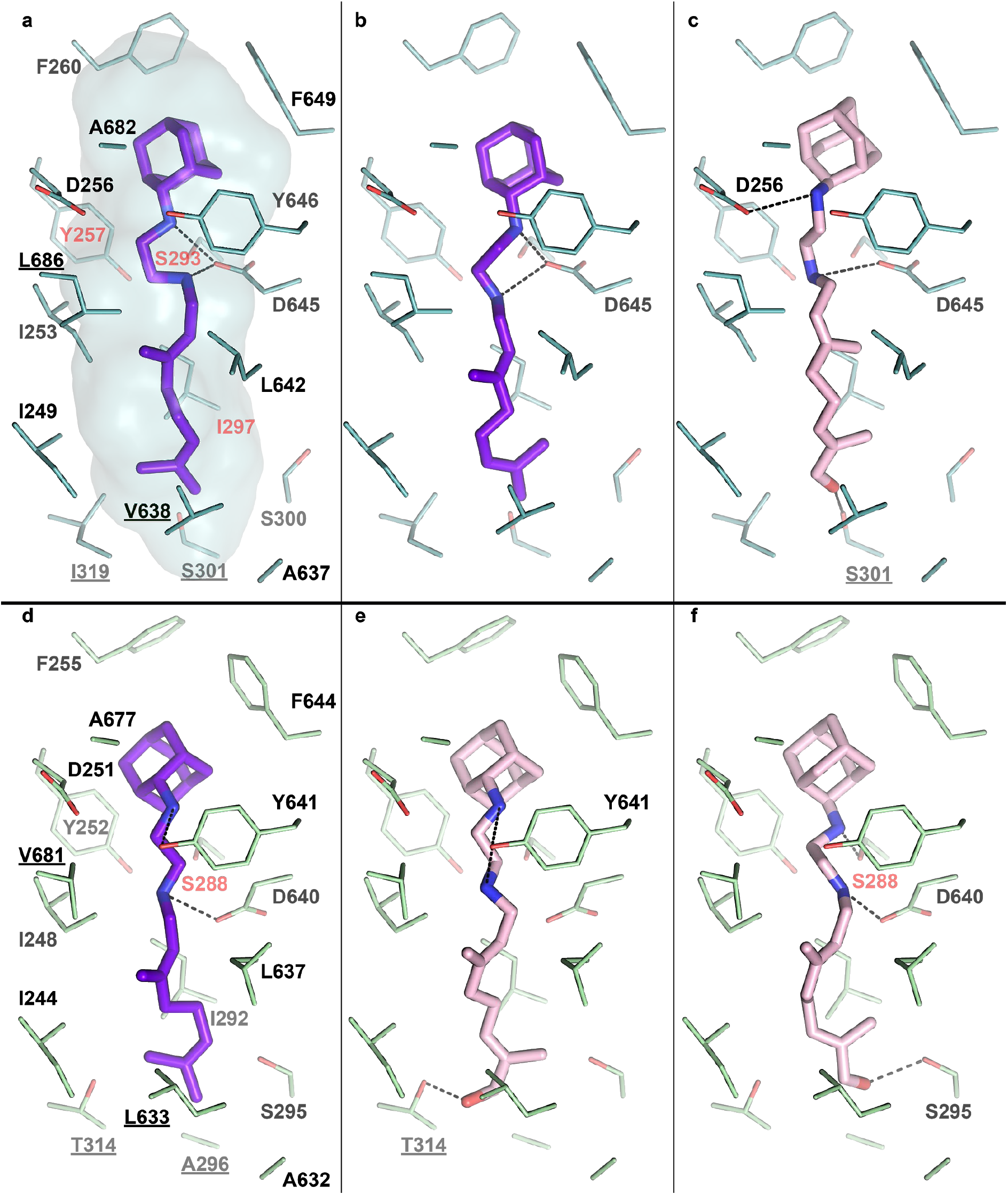
SQ109-OH docking in the Msm and Mtb MmpL3. **a**, Binding pocket in the Msm MmpL3-SQ109 crystal structure (PDB ID: 6AJG; ref. 24). **b**, Docking of SQ109 to Msm MmpL3, top-3 docking pose obtained with *AutoDock Vina* and re-scored with *Convex-PL*. **c**, Docking of SQ109-OH to Msm MmpL3, top-3 docking pose obtained with *AutoDock Vina*. **d**, Docking of SQ109 to the homologically modeled Mtb MmpL3, top-1 docking pose obtained with *AutoDock Vina* and re-scored with *Convex-PL*. **e**, Docking of SQ109-OH to Mtb MmpL3, top-7 docking pose obtained with *AutoDock Vina* and re-scored with *Convex-PL*. **f**, Docking of SQ109-OH to Mtb MmpL3, top-1 docking pose obtained with *AutoDock Vina*. Residues, different in the Mtb and Msm MmpL3 binding pockets, are underlined. Residues, mutations in which are associated with SQ109 resistance, are colored in red (Msm - Y257C, S293A/T, and I297F; Mtb - S288T; ref. 24). Possible H-bonds are shown in dotted lines.

Activation of prodrugs by Mtb enzymes was previously demonstrated for isoniazid, pyrazinamide, and ethionamide [29]. Different classes of enzymes catalyze these reactions: catalase-peroxidase encoded by katG gene [30], pyrazinamidase [31], and mycobacterial Baeyer-Villiger monooxygenases [32]. It also has been shown that Mtb acetyltransferases and phosphotransferases deactivate aminoglycosides (second-line anti-TB drugs). Xenobiotics (including anti-TB compounds) transformation in Mtb has also been shown through N–alkylation, amidation, ester hydrolysis, and the reduction of the nitro group [29]. In this work, we extend the current knowledge and demonstrate the involvement of the Mtb CYP enzyme in the hydroxylation of the anti-TB drug. Given the importance of this group of enzymes in the metabolism of xenobiotics in humans and the significant number of CYPs in Mtb, we suggest that Mtb CYPs may be involved in the metabolism of various classes of compounds. In this regard, the assessment of the anti-TB drug candidate’s metabolism using the whole cell-based system [33,34] or isolated Mtb enzymes could be a useful tool in anti-TB drug discovery.

## Materials and methods

### Cloning, expression, and purification of recombinant CYP124

cDNAs encoding CYP125 (gene Rv3545c), CYP142 (gene Rv3518c), and CYP124 (gene Rv2266) were amplified by PCR genomic DNA of Mtb H37Rv (obtained from The Vyshelessky Institute of Experimental Veterinary Medicine, NASB). Expression plasmids for each protein were generated using the vector pTrc99a. The proteins were expressed and purified as described previously [18]. The cDNA encoding spinach Ferredoxin-1 (Fdx1) was amplified from the total RNA isolated from *Spinacia oleracea* seedlings. Fdx1 was expressed in *E. coli* and purified using metal-affinity and anion-exchange chromatography.

### Substrate binding studies

To determine ligand-binding constants (Kd_app_ values) of the CYPs, optical titration was performed using a Cary 5000 UV-VIS NIR dual-beam spectrophotometer (Agilent Technologies, Santa Clara, CA) in 1-cm pathlength quartz cuvettes. Stock solutions of the steroids were prepared at a concentration of 10 mM in 45% hydroxypropyl-beta-cyclodextrin (HPCD). Titration was repeated at least three times, and Kd_app_ was calculated as described previously [18].

### Catalytic activity assay

The catalytic activity of Mtb CYPs was reconstituted in 50 mM potassium phosphate (pH 7.4) containing 0.5 μM CYP, 2 μM spinach Fdx1, 0.5 μM adrenodoxin reductase-like flavoprotein (Arh1, A18G mutant), 100 μM substrate, 1 mM glucose 6-phosphate, 1 U/ml glucose 6-phosphate dehydrogenase, and 0.4 mM β-NADPH. The proteins (CYPs, Fdx1, and Arh1) were pre-incubated with steroid substrates in the buffer solution for 10 min at 30 °C. The reaction was started by adding an NADPH-regenerating system containing glucose 6-phosphate, glucose 6-phosphate dehydrogenase, and β-NADPH. After 1 h of incubation at 30 °C, the reaction was stopped, and the reaction mixture was extracted and subjected to the LC-MS analysis. The activity of CYP124 was estimated as Turnover Number – nmoles of metabolized product / nmole of CYP / min.

### Identification of SQ109 product

An Agilent 1200 series HPLC instrument equipped with an Agilent Triple Quad 6410 mass-spectrometer was used. The samples were analyzed by gradient elution on a Zorbax Eclipse Plus C18 column (2.1 × 50 mm; 1.8 μm). TFA (0.1 % v/v in water) was used as mobile phase A and acetonitrile as mobile phase B. The gradient was 20% – 50% B in 0–7 min. The flow rate was 400 μl/min. The column temperature was maintained at 35 ± 1 °C. The mass-spectrometry experiments were performed with an electrospray ionization source (ESI) in positive-ion mode. The nebulizing gas flow rate was set at 9.5 l/min, the drying gas temperature at 350 °C, the capillary voltage at 4000 V, and the nebulizer at 35 psi. The ESI-MS/MS analysis was done in product-ion mode with different values of the fragmentor (135, 150, and 200 V) and the collision energies (10 and 20 V).

### Crystallization, data collection, and crystal structure determination

CYP124–SQ109 was crystallized by a sitting drop approach in 96-well crystallization plates with commercially available kits (Qiagen) at 20 °C with 1:1 protein/mother liquor ratio with the ligand concentration of 100 μM. Crystals were obtained from 0.3 M Mg(HCO_2_)_2_ and 0.1 M Tris pH 8.5. Glycerol (20%) as cryoprotectant was added before flash-freezing in liquid nitrogen.

The diffraction data were collected at the European Synchrotron Radiation Facility (ESRF) beamline ID23-1. The data collection strategy was optimized in *BEST* [35]. The data were processed in the *XDS* software package [36]. The crystallographic data collection statistics are given in Table 1.

**Table 1.** Crystallographic data collection and refinement statistics.

| **Data processing** |  |  |
| --- | --- | --- |
| PDB ID code |  | 6T0J |
| Source |  | ESRF ID23-1 |
| Wavelength (Å) |  | 0.972 |
| Space group |  | P12<sub>1</sub>1 |
| Cell dimensions |  |  |
|  | a, b, c (Å) | 51.54, 75.10, 56.59 |
| | $\alpha, \beta, \gamma$ (°) | 90, 106.840, 90 |
| No. of observations |  | 1056761 (63680) |
| No. of unique reflections |  | 162381 (11133) |
| Resolution (Å) |  | 30 - 1.10 (1.13 - 1.10) |
| R<sub>meas</sub> |  | 0.193 (2.751) |
| R<sub>pim</sub> |  | 0.075 (1.126) |
| I/σI | | 5.54 (0.40) |
| CC1/2 |  | 99.7 (16.8) |
| Completeness (%) |  | 97.4 (90.9) |
| Redundancy |  | 6.5 (5.7) |
| **Refinement statistics** |  |  |
| Resolution (Å) |  | 30 - 1.25 (1.28 - 1.25) |
| No. of reflections (total/unique) |  | 743229/111159 |
| R<sub>work</sub>/R<sub>free</sub> |  | 0.1479/0.1829 |
| CC* in highest shell |  | 0.816 (7601) |
| CC<sub>work</sub>/CC<sub>free</sub> in highest shell |  | 0.787/0.729 |
| No. of atoms |  |  |
|  | Protein | 3861 |
|  | Haem | 86 |
|  | SQ109 | 48 |
|  | Solvent | 830 |
| B-factors (Å<sup>2</sup>) |  |  |
|  | Protein | 14.0 |
|  | Haem | 9.7 |
|  | SQ109 | 16.6 |
|  | Solvent | 34.3 |
| R.m.s.d |  |  |
|  | Bond lengths (Å) | 0.008 |
|  | Bond angles (°) | 1.09 |
| Ramachandran statistics |  |  |
|  | Favoured (%) | 97.89 |
|  | Allowed (%) | 2.11 |

The phase problem was solved by molecular replacement in *Phaser* [37] from *PHENIX* [38], where the generated poly-ala model of substrate-free CYP124 (PDB ID code 2WM5; ref. 17) was utilized as a starting model. The space group was P12_1_1 and contained one molecule per asymmetric unit. The model was subsequently rebuilt in *PHENIX.AutoBuild* [39]. *PHENIX.Refine* [40], and *Coot* [41] were used for model refinement. In the last refinement steps, the *mF*_*o*_−*DF*_*c*_ map unambiguously showed SQ109 in the active site (Fig. 4c). The final resolution cut-off was determined by the application of paired refinement [42]. The quality of the resulting model was analyzed by *PHENIX.MolProbity* [43] and *Quality Control Check* web server.

The figures containing electron density and molecular structures were generated using *PyMOL* [44].

### Molecular docking

A 3D conformer of SQ109-OH was created in *PyMOL* [44] by adding a hydroxy group to SQ109 from its complex with Msm MmpL3 (PDB ID: 6AJG; ref. 24). Ligand torsion trees were created in *AutoDock Tools*. The structure of the Msm MmpL3 was taken from its complex with SQ109 (PDB ID: 6AJG; ref. 24). The structure of the Mtb MmpL3 was modeled using the *Phyre^2^* web server [45]. The model of the first 757 residues was based on the homology with Msm MmpL3-ICA38 complex (PDB ID: 6AJJ; ref. 24), scored with 100% confidence by *Phyre^2^*. Notably, the C-terminal domain of Mtb MmpL3 (P758-L944) was not modeled due to the absence of the alignment coverage. However, it was not used for the docking. Polar hydrogens of the protein molecules and the ligands were assigned in *PyMOL*. Molecular docking was performed, following two protocols. For the first one, we used *AutoDock Vina* [46] with default settings except for *exhaustiveness*, which was set to 10. For the second protocol, we used our in-house modification of *AutoDock Vina* and the *Convex-PL* [47] scoring function for rescoring, augmented with additional descriptors that account for conformational flexibility and solvation. We have recently applied this protocol for pose prediction in [48], where it is discussed in more detail. We ran several *AutoDock Vina* simulations to obtain more diverse docking poses and clustered all resulting conformations with a 1 Å threshold using *RDKit* [49]. To confirm the docking protocols, we compared the docked SQ109 poses with crystallographic ones (PDB ID: 6AJG; ref. 24; Fig. 4, a, b and d). The only significant difference was in the amylene part of the molecule, which may be attributed to the structure’s low resolution. We visually inspected the ten top-ranked docking poses produced by both protocols.

The figures illustrating interactions between SQ109-OH and MmpL3 proteins were generated using *PyMOL* [44] and *PLIP* [50].

## Data availability

The Structure for CYP124 in complex with SQ109 was deposited in Protein Data Bank with accession number 6T0J. The diffraction images and processing data were deposited to Integrated Resource for Reproducibility in Macromolecular Crystallography (http://proteindiffraction.org/). Other data and materials are available upon request from the corresponding authors.

## Author’s contributions

Conceptualization, A.Gi., N.S.;

Methodology, V.B., S.G., A.Gi., N.S.;

Investigation, S.B., T.V., I.G., E.M., P.S., M.K., Ant.K., A.Gu., A.Gi., N.S.;

Writing – Original Draft, S.B.;

Writing – Review & Editing, S.B., V.B., N.S., A.Gi., E.M., S.G., M.K.;

Funding Acquisition, V.B., N.S., A.Gi.,

Supervision, V.B., N.S., A.Gi.

## Acknowledgments

We thank S. Fatykhava and O. Bokut for their excellent technical assistance. We thank Prof. Rita Bernhardt (Saarland University, Saarbrucken, Germany) for providing the expression construct for Arh1. This work was supported by a joint research grant of the Belarusian Republican Foundation for Fundamental Research (B20R-061) and the Russian Foundation for Basic Research (20-54-00005). We acknowledge the ESRF Structural Biology Group and especially A. N. Popov for his assistance with the crystallographic data collection. We also thank the NIH Joint Center for Structural Genomics for the Quality Control Check web server. We are grateful to Elena Bazanova (LTTC, MIPT) for the language editing.

